# The human odorant receptor OR10A6 is tuned to the pheromone of the commensal fruit fly *Drosophila melanogaster*

**DOI:** 10.1101/2020.12.07.414714

**Authors:** Tim Frey, Charles A. Kwadha, Erika A. Wallin, Elsa Holgersson, Erik Hedenström, Björn Bohman, Marie Bengtsson, Paul G. Becher, Dietmar Krautwurst, Peter Witzgall

**Affiliations:** Leibniz-Institut für Lebensmittel-Systembiologie an der Technischen Universität München, Lise-Meitner Strasse 34, D-85354 Freising, Germany; Department of Plant Protection Biology, Swedish University of Agricultural Sciences, Box 102, 23053 Alnarp, Sweden; Department of Chemical Engineering, Mid Sweden University, Holmgatan 10, 85170 Sundsvall, Sweden; Systembolaget AB, 103 84 Stockholm, Sweden

**Keywords:** pheromone, semiochemical, food odour, aroma, odorant receptor, olfaction, Drosophila

## Abstract

**Background:** All living things speak chemical. The challenge is to discover the vocabulary, the volatile odorant chemicals that enable communication across phylogenies and to translate them to physiological, behavioural and ecological function. Olfactory receptors (ORs) interface animals with airborne odorants. Expression of single ORs in human embryonic kidney cells (HEK-293) makes it possible to interrogate ORs with synthetic chemicals and to identify cognate ligands that convey olfactory information.

**Results:** The cosmopolitan strain of the vinegar fly *Drosophila melanogaster* has accompanied the human expansion out of Africa, more than ten thousand years ago. These flies are strictly anthropophilic and depend on human resources and housing for survival, particularly in colder climate zones. Curiously, humans sense the scent of a single fly, and more precisely the female pheromone (Z)-4 undecenal (Z4-11Al), at 10 ng/mL (0.06 µmol/L). A screening of all functional human ORs in a HEK-293 assay provides an explanation for this astounding sensitivity, as it shows that OR10A6, one of the most highly expressed human ORs, is specifically tuned to Z4-11Al. Chemical analysis of fly effluvia confirms that cosmopolitan *D. melanogaster* females release Z4-11Al, while females of an African fly strain from Zimbabwe release a 1:3-blend of Z4-11Al and (Z)-4 nonenal (Z4-9Al). Interestingly, a blend of Z4-9Al and Z4-11Al produces a different aroma than the the single compounds, which is why we readily differentiate cosmopolitan and Zimbabwe flies by nose.

**Conclusion:** That we sensitively and specifically perceive the fly pheromone Z4-11Al suggests that it is a component of human odour scenes. This may have afforded a sensory drive during adaptation of commensal flies to human habitats and selected for a role of Z4-11Al in fly aggregation and premating communication. Screening ORs for key ligands leads to the discovery of messenger chemicals that enable chemical communication among and betwen vertebrate and invertebrate animals.

## Background

Volatiles from animals, microorganisms and plants are yielded by basic metabolic pathways, they inform about resources and habitats, and reveal identities and social context. Communication via volatile chemicals is reliable and inclusive, for all living beings across the kingdoms.

Volatiles are aired tweets for those equipped with receptors and sensory circuits to capture and interpret them. Animals possess olfactory receptors (ORs) for peripheral detection of volatiles, and for filtering out scents that make ecological and behavioural sense, from a noisy chemical airspace. The importance of ORs in interfacing animals with the chemical environment is reflected by the strong selection pressure on ORs to adapt to ecosystem and habitat cues and to social and sexual signals (Hayden et al. 2010, Fleischer et al. 2018, Robertson 2019, Saraiva et al. 2019). Olfaction has developed independently in invertebrates and vertebrates, but the overarching organisation of the olfactory system, building on rapidly evolving ORs, expressed in peripheral olfactory sensory neurons, feeding into a hierachy of central olfactory circuits, is rather similar in all animals (Su et al. 2009, Bear et al. 2016)

A principal, current objective and fascinating challenge in vertebrate and invertebrate olfaction research is to explore the receptive range of ORs and to set landmarks in chemical space, for a comprehension of olfactory codes and the functional analysis of olfactory systems. Expression of ORs in heterologous cell systems, for example in human embryonic kidney (HEK-293) cells, enables experimental access and makes it possible to interrogate individual ORs and to identify their cognate ligands (Krautwurst et al. 1998, Corcoran et al. 2014). ORs are seven-transmembrane domain G-protein coupled receptors (GPCRs) and a first challenge is to achieve fully functional membrane expression (Noe et al. 2017b). The ensuing step is to compose comprehensive and manageable, yet representative panels of biologically relevant compounds for investigating their receptive range. One strategy is to use key food volatiles, identified from the odorant space of food and beverages (Krautwurst & Kotthoff 2013, Dunkel et al. 2014). Odorant panels will, however, always remain notoriously incomplete - in comparison with an overwhelmingly diverse odorscape, containing countless chemicals. In vitro OR screenings do afford active ligands, but most human ORs remain in the orphan state (de March et al. 2015, Block 2018).

Chemicals we perceive at very small amounts, and which are not strictly associated with food, are particularly inspiring targets for OR screenings. One such candidate compound is (*Z*)-4-undecenal (Z4-11Al), the volatile female pheromone of the commensal fruit fly *Drosophila melanogaster*. In *Drosophila*, two isoforms of DmelOR69a, with a dual specificity for food odorants and pheromone, are co-expressed in the same OSNs. Intriguingly, we ourselves readily perceive Z4-11Al, which is released at subnanogram amounts per hour, and we reliably distinguish between the sexes, since only female flies produce this scent (Lebreton et al. 2017, Becher et al. 2018).

Cosmopolitan *D. melanogaster* flies are strictly anthropophilic, since they accompanied the human expansion from out of Africa more than 10.000 ya, and they are partially reproductively isolated from African flies, such as the Zimbabwe strain (Lachaise & Silvain 2004). The cosmopolitan fly pheromone Z4-11Al is an oxidation product of the aphrodisiac cuticular hydrocarbon (*Z,Z*)-7,11-heptacosadiene (7,11-HD) (Billeter et al. 2009, Lebreton et al. 2017). Since females from Zimbabwe produce more (*Z,Z*)-5,9-heptacosadiene (5,9-HD) than 7,11-HD (Dallerac et al. 2000, Grillet et al. 2012), it follows that these flies would release another aldehyde. Posing that our perception of Z4-11Al is not only sensitive but also specific, we asked whether we are able to olfactorily discriminate between females of the cosmopolitan and Zimbabwe strains of *D. melanogaster*. A sensory panel corroborated this idea by comparing synthetic compounds and fly odours.

Naturally, this begs the question - how do humans smell the scent of the fly? A range of human ORs is tuned to straight-chain aldehydes, which are commonly found in fruit and vegetable aroma (Schmiedeberg et al. 2007, Saraiva et al. 2009, Nara et al. 2011, Li et al. 2014, de March et al. 2015, Block et al. 2018) and perception of Z4-11Al might be encoded by one or even several of these aldehyde-responsive ORs. We hence submitted Z4-11Al to an in vitro screening of all functional human ORs, using heterologous expression in human embryonic kidney cells (HEK-293) and a luminescence-based assay (Noe et al. 2017a,b). This screening renders OR10A6 as the sole responsive OR for Z4-11Al. A subsequent olfactory panel test confirmed the results of an in vitro dose-response test of several synthetic aldehyde analogues, showing that we discriminate between structurally related aldehydes and that our olfactory perception of Z4-11Al is remarkably sensitive and specific.

The scent of the fly illustrates how chemical ecology research inspires the discovery of OR ligands and provides another account for chemical communication across phylogenies.

## Materials and methods

### Chemicals

Isomeric purity of (*Z*)-4-undecenal (Z4-11Al) was 98.6%, according to gas chromatography coupled to mass spectrometry (6890 GC and 5975 MS, Agilent Technologies, Santa Clara, CA, USA). Isomeric purity of (*Z*)-4-nonenal (Z4-9Al) and (*Z*)-6-undecenal (Z6-11Al) were 97.4% and 96%, repectively. Chemical purity of these synthetic aldehydes was >99.9%. Ethanol (redistilled; Merck, Darmstadt, Germany) was used as solvent.

For the OR screening assays, the following chemicals were used: Dulbecco’s MEM medium (#F0435), FBS superior (#S0615), L-glutamine (#K0282), penicillin (10000 U/ml)/streptomycin (10000 µg/ml) (#A2212), trypsin/EDTA solution (#L2143) (Biochrom, Berlin, Germany), CaCl2*2H2O (#22322.295), D-glucose (#101174Y), dimethyl sulfoxide (DMSO) (#83673.230), HEPES (#441476L), potassium chloride (#26764.230), and sodium hydroxide (#28244.295) (VWR Chemicals BDH Prolabo, Leuven, Belgium), sodium chloride (#1064041000, Merck, Darmstadt, Germany), ViaFect™ Transfection Reagent (#E4981, Promega, Walldorf, Germany), D-luciferin (beetle) monosodium salt (#E464X, Promega, Walldorf, Germany), Pluronic® PE 10500 (#500053867, BASF, Ludwigshafen, Germany), (R)-(-)-carvone (#W224908, Sigma-Aldrich, Steinheim, Germany).

### Insects

Cosmopolitan (Dalby) and Zimbabwe (S-29, Bloomington) strains of *D. melanogaster* were reared on a standard sugar-yeast-cornmeal diet at room temperature (25±2°C) and 50±5 rH under a 12:12-h L:D photoperiod. Eclosing flies were collected every 4 h and sexed under CO_2_, according to the sex comb on the third segment of the male forelegs. Presence of meconium was used as a distinguishing feature for virgin flies. Females were kept separately in 30-mL Plexiglas vials with fresh food.

### Pheromone collection and chemical analysis

Sixty unmated cosmopolitan and Zimbabwe females, respectively (n=9 and n=10, respectively) were transferred to standard glass rearing vial (24.5 x 95 mm, borosilicate glass; Fisher Scientific, Sweden), which had been baked at 350°C overnight. After 24 h, the flies were removed and the vial was rinsed with 200 µl of hexane, containing 100 ng decanal as internal standard, in an ultrasonic water bath for 3 min. The solvent was transferred to 1.5 mL GC-MS vials with insert and condensed to ca. 5 µl in a fume hood.

Two µl of the solvent rinses were analyzed by gas chromatography-mass spectrometry (GC-MS) (6890 GC and 5975 MS, Agilent, Santa Clara, CA, USA) on a fused silica capillary column (60 m x 0.25 mm), coated with HP-5MS UI (d_f_ =0.25 µm; Agilent). Injections were made in splitless mode (30 s), at 275°C injector temperature. The GC oven was programmed from 50 to 250°C at 8°C/min (2 min and 10 min hold, respectively) and a final temperature of 275°C, the mobile phase was helium (34 cm/s). The MS operated in scanning mode. Aldehydes were identified according to m/z spectra and Kovats retention indices, using custom and NIST libraries, in comparison with synthetic standards.

### OR expression and screening

#### Cell culture and transient DNA transfection

Human embryonic kidney (HEK-293) cells were cultivated in Dulbecco’s MEM medium (DMEM: w 3.7 g/L NaHCO3, w 4.5 g/L D-glucose, w/o L-glutamine, w/o Na-pyruvate) supplemented with 10% fetal bovine serum (FBS superior), 2 mM L-glutamine, 100 units/mL penicillin, and 100 µg/mL streptomycin in 10 cm cell culture dishes at 37 °C, 5% CO2, and 100% humidity, as test cell systems for the functional expression of recombinant ORs as described previously (Geithe et al. 2015, 2017b; Noe et al. 2017a). One day before transfection, HEK-293 cells were transferred with a density of 12000 cells per well in white 96-well plates (Thermo Scientific™ Nunc™ F96 MicroWell™, white, #136102, Thermo Fisher Scientific, Waltham, USA). The transfection was done by the cationic lipid-transfection method using 100 ng OR plasmid-DNA, 50 ng olfactory G-protein Gαolf (Jones & Reed 1989, Shirokova et al. 2005), 50 ng RTP1S (Saito et al., 2004), 50 ng Gγ13 (Li et al. 2013), and 50 ng genetically modified cAMP-luciferase pGloSensor™-22F (Binkowski et al. 2009) (Promega, Madison, USA), each with ViaFect™ Transfection Reagent. As negative control, transfection of an empty pFN210A-vector-plasmid (mock) was employed. As positive control OR1A1 was transfected on each plate. Each transfection was done in triplicates on the same 96-well plate. The cells were taken into experiment 42 h post-transfection as reported (Geithe et al. 2015, 2017b; Noe et al. 2017a).

#### cAMP luminescence assay

Cell culture media of the transfected HEK-293 cells in the 96-well plates was replaced 1 h prior to the luminescence measurement with physiological salt solution containing 140 mmol/L NaCl, 10 mmol/L HEPES, 5 mmol/L KCl, 1 mmol/L CaCl2, 10 mmol/L D-glucose and 2% D-luciferin, pH 7.4. After this incubation, basal luminescence signals for each well (three consecutive data points, 60 s intervals) were recorded with the GloMax® Discover Microplate Reader (Promega, Madison, USA) before odorant application. As positive control, 30 µmol/L (R)-(-)-Carvone was applied on the OR1A1 transfected cells. Odorant stock solutions were prepared in DMSO, and diluted 1:1000 into the physiological salt solution containing 0.05% Pluronic® PE 10500, as solvent mediator. Final DMSO concentration on the cells was 0.1%. 4 min after odorants were applied to the cells, three consecutive data points at 60 s intervals were recorded for each well with the GloMax® Discover Microplate Reader.

#### Data analysis of cAMP luminescence measurements

The raw luminescence data obtained from the GloMax^®^ Discover Microplate Reader were processed as followed. For each well, the average of the three data points before odorant addition was subtracted from the average of the three data points after stimulation (Δsignal). Then, the corresponding mock of each substance/concentration was subtracted from each Δsignal value. All mock-subtracted Δsignal-value were then normalized to the positive control of each plate (screening experiments), or to the respective maximum signal of each concentration-response relation. EC_50_ values were obtained by fitting the function f = ((a-d)/(1+(×/EC50)^h)+d) to the data.

### Sensory evaluation

Synthetic compounds were diluted in redistilled ethanol (Sigma-Aldrich). One h prior to testing, 10-µL aliquots were pipetted into 20-mL screw-top glass vials (Genetec) containing 10 mL of redistilled water or 10 mL of white wine (Ruppertsberger Riesling, Systembolaget, 72038-01). For evaluation of fly odour, 5 live *D. melanogaster* females, of the cosmopolitan (Dalby) and Zimbabwe strain (S-29, Bloomington), respectively, were placed during 3 h in 20-mL vials, they were released ca 30 min before testing, and 10 mL of redistilled water or wine was added to the vial.

A first pairwise comparison comprised vials containing either 10 ng synthetic Z4-11Al or fly odour, in 10 mL water or wine. Vials were assigned a random letter. Panelists (n=21) were asked whether or not the odours in the vials bear resemblance to each other. Results were compared using a paired Student’s t-test.

Subsequent triangle tests included (a) vials exposed to cosmopolitan and Zimbabwe *D. melanogaster* female flies (n=45 judges), (b) 10 ng Z4-9Al or 10 ng Z4-11Al (n=45), (c) a blend of 10 ng Z4-9Al and 3 ng Z4-11Al (n=41), and (d) and a dose-response test, of 6 different amounts of synthetic Z4-9Al, Z4-11Al and Z6-11Al, from 0.01 ng to 10 ng (n=9), in 10 mL water. Of three stimuli in each triangle, two were the same and judges were asked to point out the odd sample. Sample order and lettering of vials were randomized, a Chi2-test was used for statistical comparison.

## Results

### Sensory evaluation of fly odour vs Z4-11Al

A sensory panel compared synthetic Z4-11Al with the scent of *D. melanogaster* females, of the cosmopolitan and the Zimbabwe strains, in water and wine, providing a rich odorant background (Figure 1).

**Figure 1.**
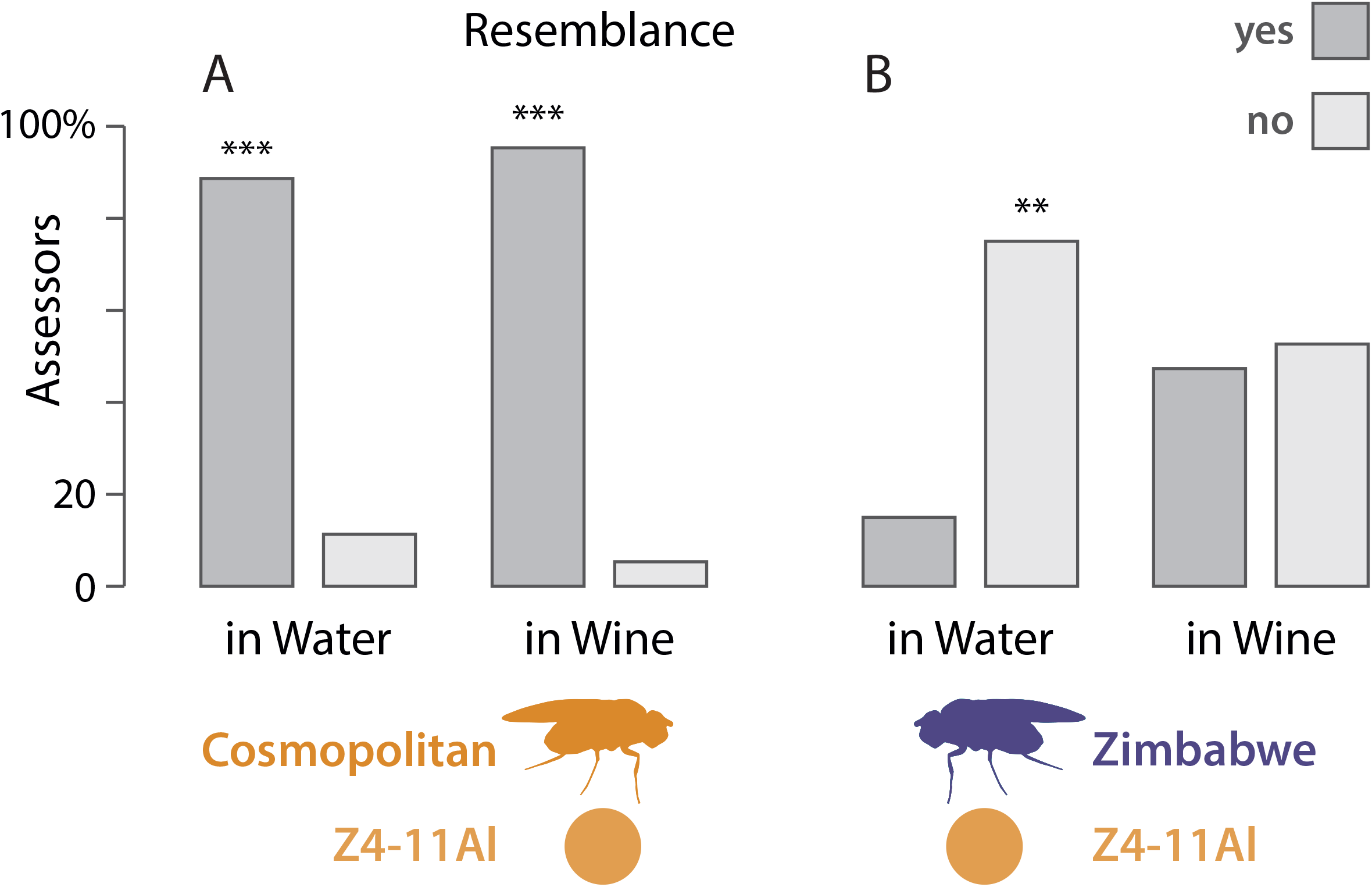
Comparison of the olfactory resemblance of 10 ng synthetic Z4-11Al and the odour of *D. melanogaster* female flies, of the cosmopolitan (A) and Zimbabwe strain (B), in water and wine. Judges were asked whether or not the odours of two vials bear resemblance. Bars marked with asterisks are significantly different (Students t-test).

Vials, where 5 fly females had been kept during 3 h and released 30 min before testing, and vials formulated with 10 ng Z4-11Al, were filled with 10 mL water or wine, respectively. In vials filled with water, a significant number of panelists found the odour of Z4-11Al to resemble cosmopolitan flies (P=0.0002), but not Zimbabwe flies (P=0.002; Figure 1). In wine, 19 out of 21 panelists readily perceived cosmopolitan fly odour and found it to resemble synthetic Z4-11Al (P<0.0001). In contrast, the evaluation of Zimbabwe fly odour vs. Z4-11Al was impaired in wine. The number of panelists who distinguished and did not distinguish Z4-11Al from Zimbabwe flies was not different at P>0.05 (Figure 1).

### Aldehyde emission by cosmopolitan and African flies

Cosmopolitan *D. melanogaster* females produce the courtship pheromone 7,11-HD, which affords Z4-11Al as oxidation product (Billeter et al. 2009, Lebreton et al. 2017). Females of the Zimbabwe strain produce mainly 5,9-HD instead (Dallerac et al. 2000), and it follows that these flies would therefore also release Z4-9Al (Figure 2A).

**Figure 2.**
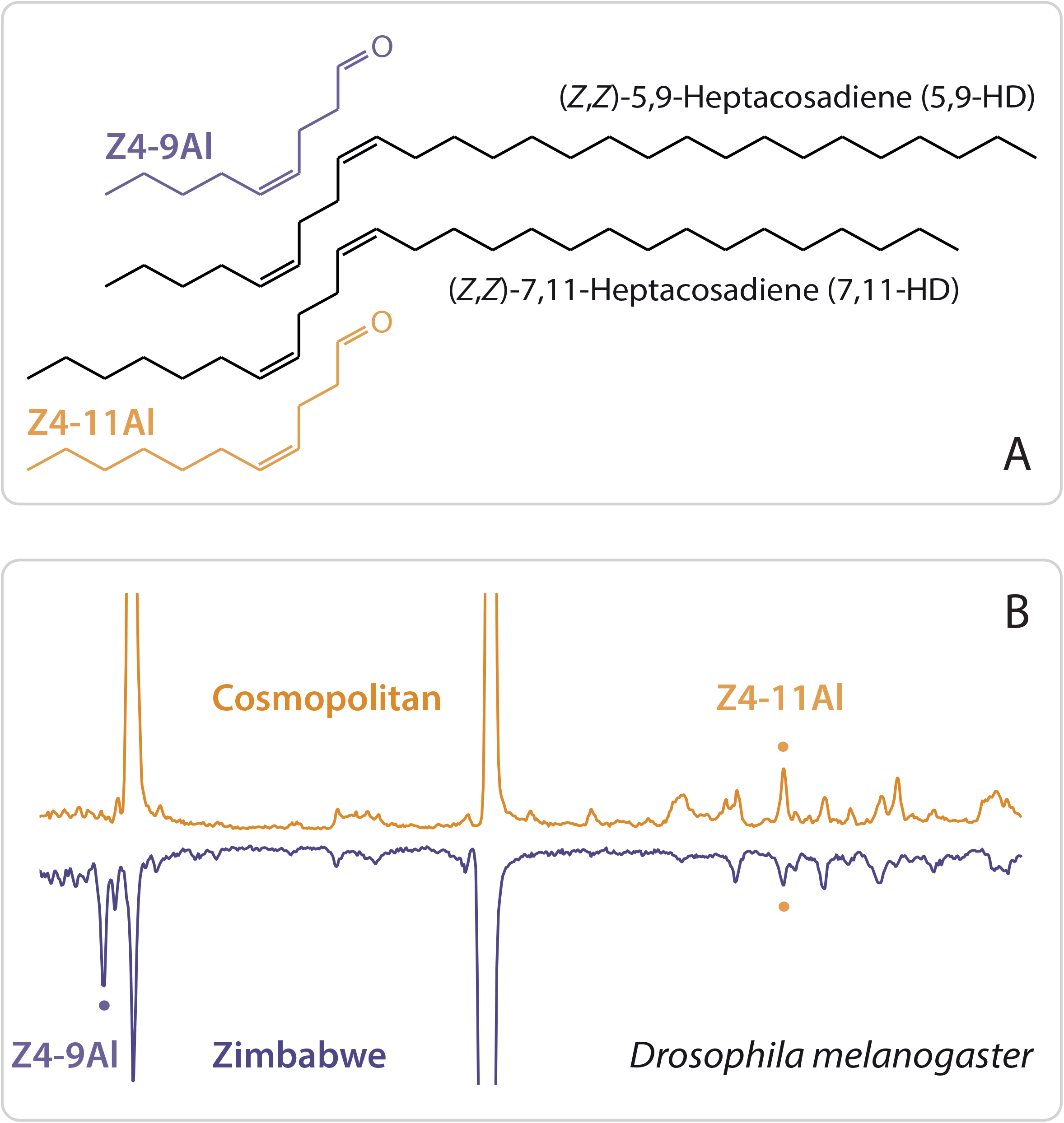
(A) The female pheromone of cosmopolitan *D. melanogaster* (*Z*)-4-undecenal (Z4-11Al) and its precursor, (*Z,Z*)-7,11-heptacosadiene (7,11-HD) (Lebreton et al. 2017). Females of the Zimbabwe strain produce in addition (*Z,Z*)-5,9-heptacosadiene (5,9-HD) and the corresponding oxidation product is (*Z*)-9-nonenal (Z4-9Al). (B) Chromatograms of headspace collections from cosmopolitan and Zimbabwe females, with peaks of Z4-9Al and Z4-11Al highlighted. Zimbabwe flies produce Z4-9Al in a 2.6±0.7-fold amount (n=10), compared to Z4-11Al.

Headspace analysis of *D. melanogaster* confirmed that females of the cosmopolitan strain released Z4-11Al, while Z4-9Al remained below detection level in all fly effluvia collections. Zimbabwe females, on the other hand, produced Z4-9Al, in addition to Z4-11Al, in a 2.6±0.7:1 ratio amount (n=10) (Figure 2B).

### The human olfactory receptor OR10A6 is tuned to Z4-11Al

OR10A6 L_287_P showed by far the strongest response to Z4-11Al, in a screening of 579 human ORs expressed in human embryonic kidney cells (HEK-293) (Figure 3). A dose-response assay confirmed that Z4-11Al was a most active ligand for OR10A6 L_287_P, compared with the aldehyde analogue Z4-9Al and the positional isomer Z6-11Al (Figure 4A).

**Figure 3.**
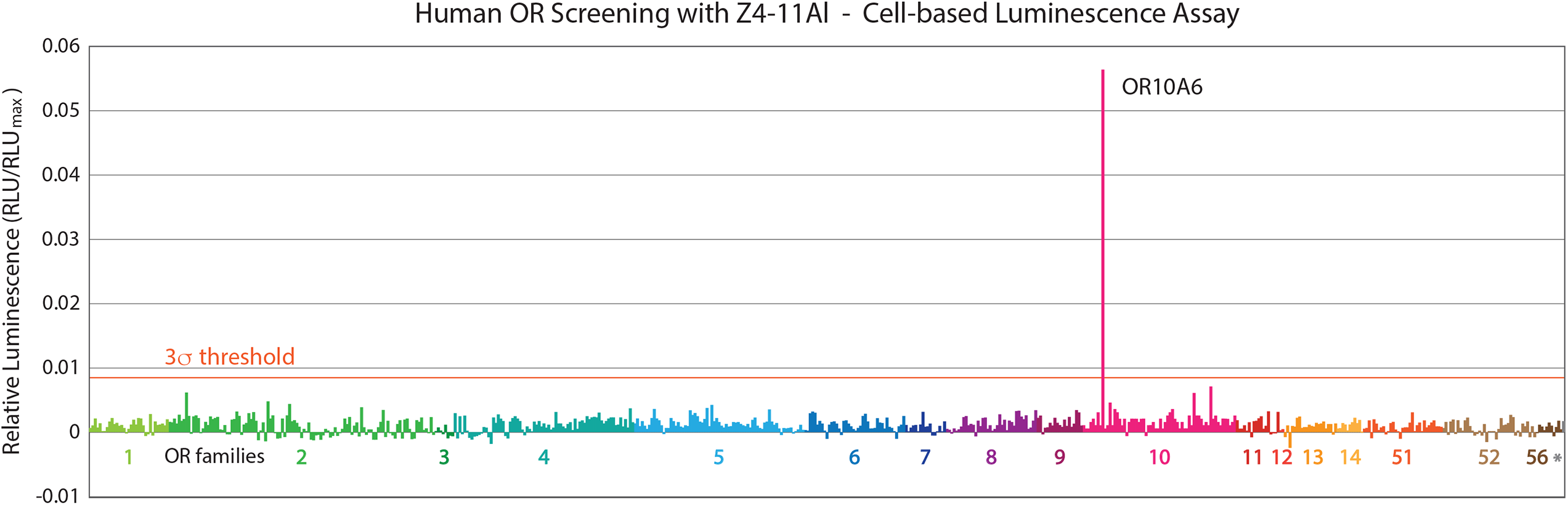
Screening of Z4-11Al (100 µmol/L) against 579 OR variants, including all 390 human OR reference sequences (according to NCBI) and their most frequent variants, showed that soleley OR10A6 L_287_P responded to Z4-11Al. Shown are means (n=2), mock control was subtracted. Relative luminescence units (RLU), normalized to the response of OR1A1 to (R)-(−)-carvone (30 µmol/L) (RLUmax).

**Figure 4.**
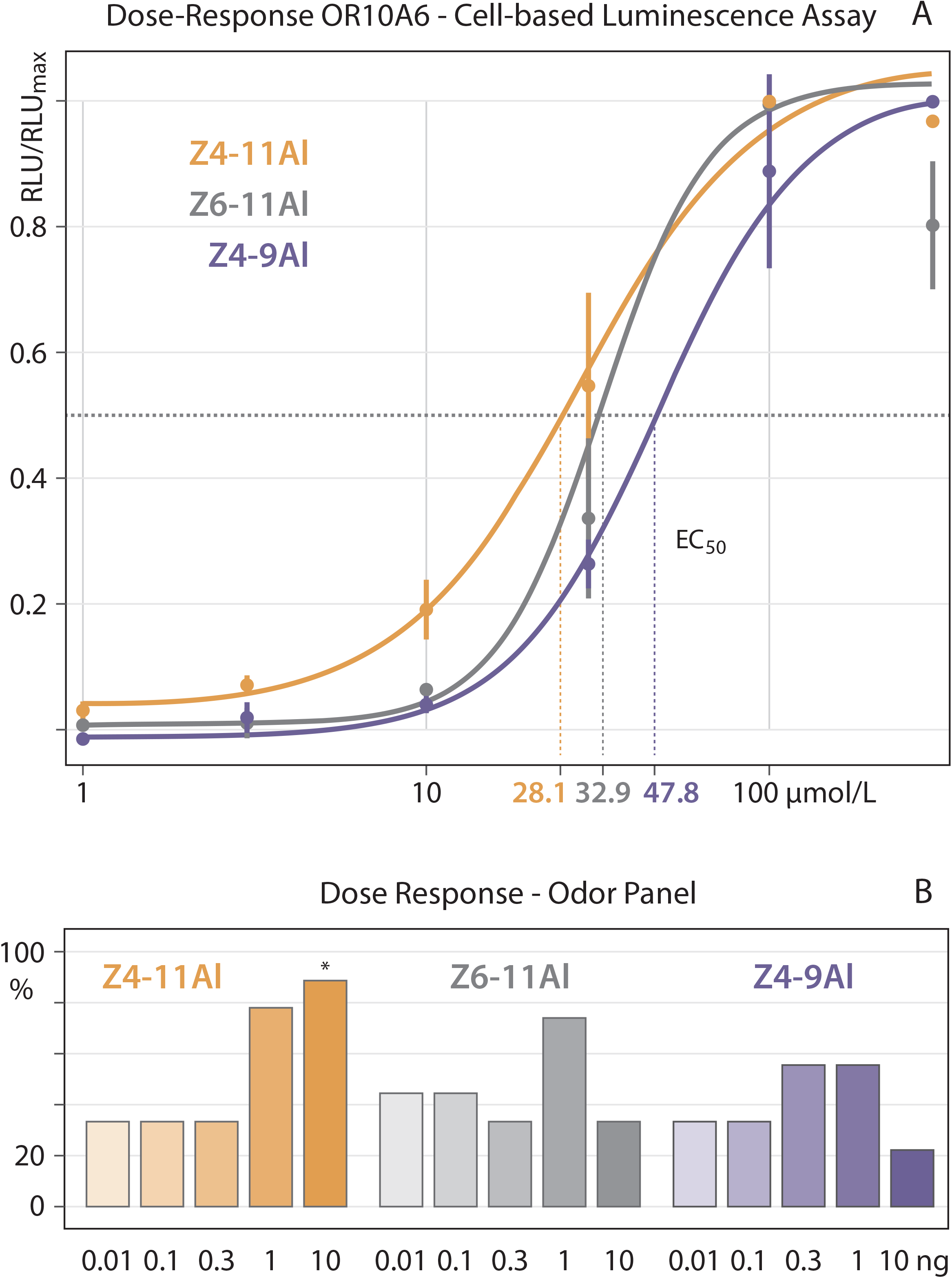
Dose response effect of synthetic Z4-11Al, Z4-9Al and Z6-11Al on OR10A6 L_287_P in HEK-293 cells (A) and an odorant panel (B). Relative luminescence units (RLU) were mock control-subtracted, and normalized to the highest Z4-11Al-induced signal of OR10A6 L_287_P (mean ± SD; n=3). For the panel test, compounds were formulated, from 0.01 ng to 10 ng, in 10 mL redistilled water. For each dose, judges (n=9) evaluated a triangle of two identical and one odd sample, they were asked to pick the odd sample. A significant number of panelists distinguished Z4-11Al at 10 ng (Chi2-test; p<0.02).

The odorant panel corroborated that we are more sensitive to Z4-11Al than to Z4-9Al or Z6-11Al (Figure 4B). A significant number of panelists sensed Z4-11Al at 10 ng/mL in water (0.06 µmol/L). In comparison, panelists did not perceive Z4-9Al or Z6-11Al, at the amounts tested.

A low response threshold to Z4-11Al, in vitro (Figure 4A) and in vivo (Figure 4B) corroborates our remarkable sensitivity to the female pheromone of cosmopolitan *D. melanogaster*. Panelists who discriminated Z4-11Al from control at the higest dose (Figure 4B) associated its aroma with citrus or grapefruit. In comparison, the aroma of Z4-10Al is reminiscent of tangerine (Douglas et al. 2001).

### Specific perception of Z4-11Al

In a subsequent triangle test, panelists discriminated between cosmopolitan and Zimbabwe female flies (P=0.01; Figure 5A). Discrimination between Z4-9Al and Z4-11Al (P=0.02; Figure 5B), however, cannot account for this observation. Emission of a blend of Z4-11Al and Z4-9Al, by Zimbabwe strain females (Figure 2), together with our far higher sensitivity to Z4-11Al than to Z4-9Al (Figure 4), seemingly contradicts a sensory differentiation between Z4-11Al and Zimbabwe flies (Figure 1) and between the two fly strains (Figure 5A).

**Figure 5.**
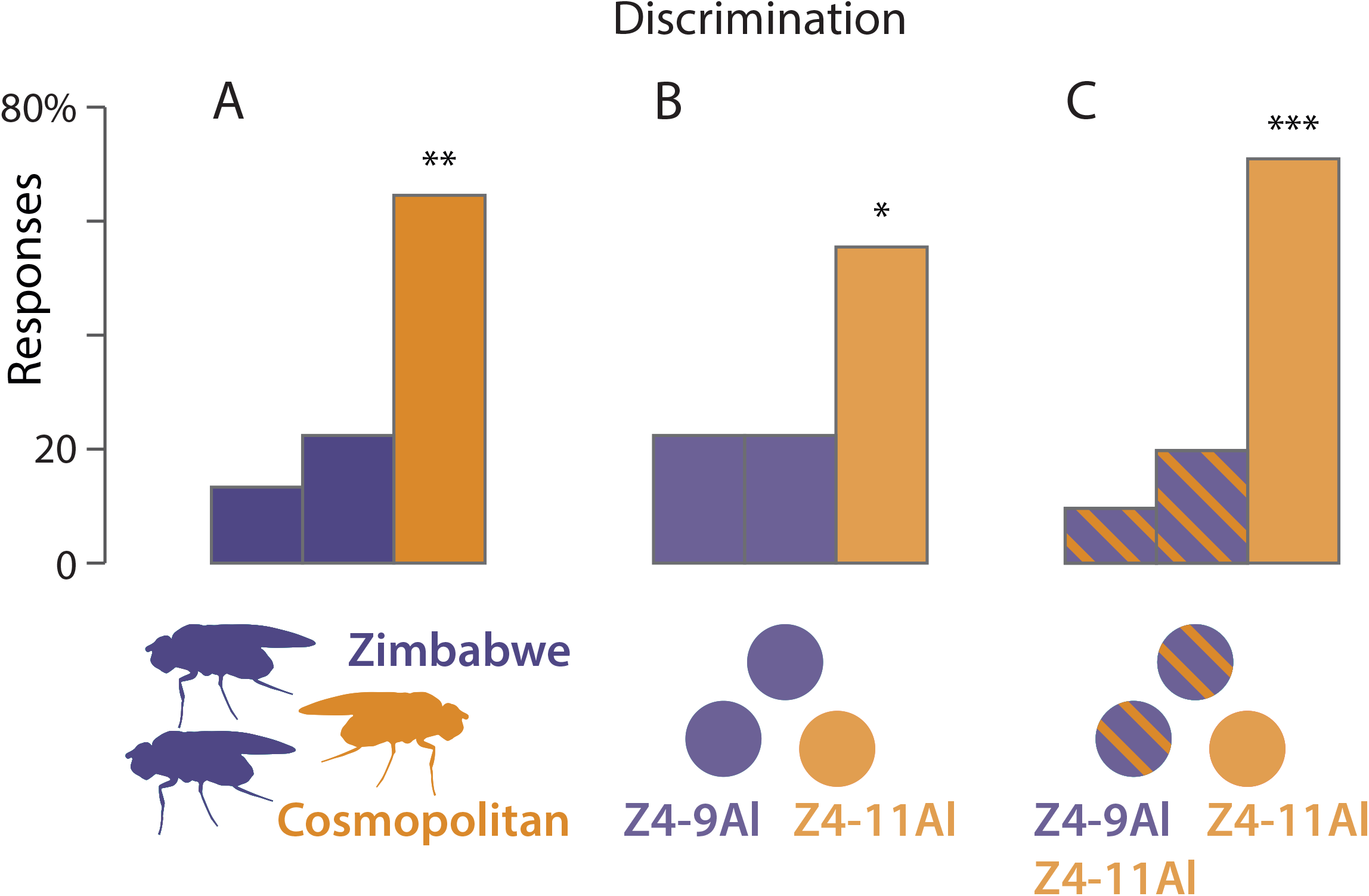
Olfactory discrimination of (A) cosmopolitan vs Zimbabwe *D. melanogaster* females, (B) 10 ng Z4-9Al vs 10 ng Z4-11Al, and (C) a 10:3-ng blend of Z4-9Al and Z4-11Al vs 10 ng Z4-11Al. Judges were asked to point out the odd sample, among three vials containing two identical samples. Bars marked with asterisks are significantly different (Chi2-test). Vials containing synthetic compounds and fly odour were filled in 10 mL redistilled water.

Cosmopolitan and Zimbabwe female flies are expected to differ only with respect to odour intensity, not quality - unless a blend of Z4-11Al and Z4-9Al produces a different aroma than the Z4-11Al. This is indeed the case. A triangle test shows a very clear distinction between Z4-11Al and a 3:10-blend of Z4-11Al and Z4-9Al (P=0.01; Figure 5C) mimicking the scent of cosmopolitan and Zimbabwe flies, respectively.

## Discussion

### Z4-11Al is a key ligand for the highly expressed OR10A6

Our olfactory perception of Z4-11Al, the female pheromone of the cosmopolitan strain of *D. melanogaster* is highly sensitive and specific. We sense synthetic Z4-11Al at 10 ng/mL (0.06 µmol/L) (Figure 4B) or at subnanogram amounts released by single flies (Figure 1; Lebreton et al. 2017), and we distinguish between Z4-11Al and structurally similar aldehydes, such as Z4-9Al and Z6-14Al (Figure 4B), and even a blend of Z4-11Al and Z4-9Al (Figure 5C).

We here show that OR10A6 L_287_P (OR10A6), a very highly expressed human olfactory receptor (Saraiva et al. 2019), is specifically tuned to Z4-11Al (Figure 3). Other human ORs known to respond to odour-active aldehydes that are, for example, part of fruit aromas (Nara et al. 2011, de March et al. 2015, Block 2018) are not nearly as responsive to Z4-11Al as OR10A6 (Figure 3).

Remarkably, the panel also discriminated between Z4-11Al and a blend of Z4-11Al and Z4-9Al, which explains how we discriminate cosmopolitan from Zimbawe flies (Figure 2, 5). Since OR10A6 is responsive to Z4-9Al (Figure 4), it is possible that modulation at the OR level encodes this blend discrimination. Synergistic and antagonistic responses in olfactory neurons to odour mixtures, compared with the individual components, show that processing of odour interactions is not restricted to higher olfactory circuits, but occurs even peripherally (Brann & Datta 2020, McClintock et al. 2020, Xu et al. 2020).

Broadly tuned, yet highly selective ORs are the basis for sensing a diverse odorant environment with only a limited number of ORs (Geithe et al. 2017a, Block 2018, Saraiva et al. 2019). Odorant interaction and encoding of odorant mixtures at the OR level very substantially extends the receptive range of ORs.

It is further intriguing that we sense Z4-11Al released by single flies against the rich bouquet emerging from a glass of wine (Figure 1A; Becher et al. 2018). Single ORs and their key ligands play indeed a central role in olfactory object recognition, especially against heterogenous backgrounds. Olfactory sensory neurons expressing high-affinity ORs with low activation thresholds have been shown to become activated early during a sniff and thus accentuate the response to behaviourally salient signals, while input from other ORs is temporarily tuned down (Wilson et al. 2017, Arneodo et al. 2018, Bolding & Franks 2018).

Taken together, our observations illustrate how a single OR contributes to olfactory perception, in addition to combinatorial coding of odorant blends by arrays of several ORs (Mainland et al. 2014). Sensitivity to odorants also depends on the number of olfactory sensory neurons expressing the corresponding ORs (van der Linden et al. 2020). OR10A6 is among the most highly expressed ORs in our nose (Verbeugt et al. 2014, Saraiva et al. 2019), which likely accounts for our high sensitivity to Z4-11Al.

### Occurrence of Z4-11Al in human odour scenes

Highly abundant ORs are plausibly dedicated to odorants of critical physiological, behavioral or ecological function, and this raises the question what Z4-11Al means to us. Perception of the same odorant by insects and vertebrates is convergent, since the respective ORs are built differently and lack a common phylogenetic root (Su et al. 2009, Bear et al. 2016). It is, however, unclear whether shared perception is merely coincident, or whether Z4-11Al is of mutual importance for flies and men.

What is the source of Z4-11Al in a human odorscape? Animals, plants and associated microbes each release many hundreds of compounds and these volatile emissions change with age, phenology and phyisological state (Knudsen et al. 1993, El-Sayed 2020, Lemfack et al. 2018, Ljunggren et al. 2019). Z4-11Al has not been searched for, synthetic standards are not commercially available and we can safely assume that the occurrence of Z4-11Al is only incompletely known.

In food plants, Z4-11Al has been found in coriander and clementine (Chisholm et al. 2003, Eyres et al. 2005), and similar aldehydes are typical for fruit (e.g. Fischer et al. 2008, Chai et al. 2012). Monoenic aldehydes are perceived as “citrusy”, but also as “tallowy” since they contribute to the flavour of cooked or roast food and meat, including rice, oils, fish, chicken and beef (Cha et al. 1992, Siegmund and Pfannhauser 1999, Rochat and Chaintreau 2005, Roh et al. 2006, Yang et al. 2008, Oueslati et al. 2018), and Z4-11Al has also been found in oxidized tallow (Shi et al. 2013).

The crested auklet, a sea bird, releases a tangerine-scented, social odour that signals mate quality and contains (*Z*)-4-decenal as a main compound (Douglas et al. 2001, Hagelin et al. 2003). Unsaturated aldehydes are part of human skin emanations (Duffy et al. 2018) and serve as cues for detecting the presence of humans (Li et al. 2014). Intriguingly, milk from humans and rabbits contains (*E*)-2-decenal and (*Z*)-2-undecenal, respectively, and Z4-11Al has been found in rabbit anal glands, accelerating heartbeat upon perception (Goodrich et al. 1978, Schaal et al. 2003, Duffy et al. 2018). Taken together, Z4-11Al is found in human food, it might even be produced by ourselves and could manifest food, social context, or both.

### Emission of Z4-11Al by *D. melanogaster*

The vinegar fly is probably our first, involuntarily domesticated animal and it accompanied our global expansion from out of Africa more than ten thousand years ago. Cosmopolitan vinegar flies are associated with us on all continents and most climate zones, they are strictly anthropophilic, depend on our dwellings and food for survival and they share our taste for fermenting food (Lachaise & Silvain 2004, Arguello et al. 2019). *D. melanogaster* females, not males, produce dienic hydrocarbons that give rise to monoenic aldehydes, which is why we smell the female flies (Everaerts et al. 2010, Lebreton et al. 2017).

The sibling species *D. simulans* has also attained worldwide distribution in association with humans, but is, unlike *D. melanogaster*, not a strict commensal and more rarely found in households or buildings (Lachaise & Silvain 2004). *D. simulans* females do not produce dienic hydrocarbons, which is a main element of the mating barrier between these two sibling species. The cuticular hydrocarbon 7,11-HD promotes courtship in cosmopolitan *D. melanogaster*, and suppresses interspecific matings with *D. simulans*, owing to differential, species-specific coding of 7,11-HD in neural circuits mediating reproductive behaviour (Billeter et al. 2009, Billeter & Wolfner 2018, Seeholzer et al. 2018, Sato and Yamamoto 2020).

Cosmopolitan and African *D. melanogaster* females also differ with respect to cuticular hydrocarbons. The female-specific desaturase gene desat2, which affords 5,9-HD and Z4-9Al, is functional only in African and not in cosmopolitan flies (Figure 2; Dallerac et al. 2000, Grillet et al. 2012). This hydrocarbon polymorphism yields a distinctive aldehyde blend, which which is how we differentiate the scent of these two fly strains (Figure 5).

Species-specific differences in hydrocarbons align with corresponding aldehyde signatures, that entail behavioural consequences in the flies. Z4-11Al attracts *D. melanogaster*, but not males of the Zimbabwe strain, and has an antagonistic effect on upwind flight attraction in *D. simulans*. This underlines the role of female-produced volatile pheromones in long-range mate communication in *Drosophila* (Lebreton et al. 2017).

### Chemosensory habitat adaptation and sensory drive

ORs are known to readily adapt to habitats and to dietary or social chemosensory niches, in insects and vertebrates alike (Bear et al. 2016, Hughes et al. 2018, Saraiva et al. 2019). Splice forms of the fly receptor DmelOR69a are tuned to food odorants, including yeast and citrus fruit flavours, and to the female pheromone, respectively (Lebreton et al. 2017). These splice forms provide an extra degree of freedom for acquisition of or adaptation to new ligands, as long as they match the food and mate finding theme.

We are food and home to the flies, they depend on us and especially so in cold climate. If Z4-11Al is part of human odour scenes, it provides a suitable signal for presence of humans and related food resources, especially since it already participates in mate communication in African *D. melanogaster*. Adaptation to a commensal lifestyle may have provided a sensory drive and selection pressure towards a single pheromone, Z4-11Al.

The interaction between natural and sexual selection has indeed ben shown to effect cuticular hydrocarbon composition and mate recognition in *D. melanogaster* (Blows 2002). Habitat selection and specific mate recognition are tightly interconnected (Paterson 1985, Endler 1992, Boughman 2002) and a chemosensory bias for an odorant that is characteristic for human habitats may have driven a role of such an odorant in sexual premating communication.

## Conclusion

Sensing the scent of a single fly is out of the ordinary (Becher et al. 2018), and we even differentiate between cosmopolitan and Zimbabwe female pheromones (Figure 5). But only the discovery that a single, most highly expressed human OR (Figure 3; Saraiva et al. 2019) is tuned to the fly pheromone Z4-11Al underlines the biological significance of these observations.

Sensitive and specific perception suggests that Z4-11Al is found in human habitats, which may have driven the role of Z4-11Al in mate communication in commensal *D. melanogaster* flies. Convergent perception of Z4-11Al may stem from shared food resources and associated microorganisms. This is reminiscent of dedicated olfactory channels for geosmin that alert flies and humans about the presence of mould, which is detrimental for all animals alike (Maga 1987, Stensmyr et al. 2012).

Ambient odorscapes contain countless chemicals of yet unknown activity. Our study highlights how the identification of key OR ligands leads to the discovery of salient messenger chemicals and delivers insights in how chemical communication interconnects species across phylogenies. Regrettably, we can barely speculate what the fly pheromone may mean to us and whether it signals food, social context, or both. Satisfying our curiosity is an excellent reason to pursue, since the vinegar fly continues to afford fundamental discoveries and studying fly sex perfumes may perhaps teach us about our own.

### Availability of data and materials

Data will be uploaded to Dryad.

## Funding

This study was fincially supported by the Faculty of Landscape Architecture, Horticulture, and Crop Production Science (SLU, Alnarp, Sweden).

## Authors’ contributions

PW, DK and PGB conceived the study. OR-screening by TF and DK, fly chemical analysis by CAK, BB and MB. Sensory panel tests by EH, PGB and PW. EAW and EH synthesized aldehydes. PW wrote text with input from all co-authors, all authors read and approved the final manuscript version.

## Acknowledgements

We thank the members of the sensory panel (Systembolaget, Stockholm) for evaluating the fly scent.

## Ethics approval and consent to participate

Not applicable.

## Consent for publication

Not applicable.

## Competing interests

Authors declare no competing interests.

## Notes

### Competing Interest Statement

The authors have declared no competing interest.

